# Transcriptomic Signatures of Neuroinflammation and Adaptive Immunity in the Human SCG During Cardiac Disease: A Bulk RNA-seq Reanalysis

**DOI:** 10.1101/2025.05.18.654713

**Authors:** Nourelden Rihan

**Affiliations:** Medical Student, Faculty of Medicine, October 6 University, Cairo, Egypt

**Keywords:** Sympathetic nervous system, Superior cervical ganglia, Cardiac disease, RNA sequencing, Neuroinflammation, Immune-metabolic signaling, Human transcriptomics

## Abstract

The superior cervical ganglia (SCG) play an important role in sympathetic and autonomic regulation, yet their molecular adaptations in cardiac disease remain poorly understood. Here, we present an independent exploratory reanalysis of publicly available human bulk RNA-seq data from SCG samples of cardiac disease patients (n = 3) and healthy controls (n = 3) (ENA: PRJNA967653) to uncover novel transcriptomic insights. Differential expression and functional enrichment analyses (GO, KEGG, GSEA) revealed significant neuroinflammatory signatures in cardiac disease, including upregulation of inflammatory chemokines, macrophage activation markers, and complement system components, alongside enrichment of adaptive immune processes, marked by increased expression of multiple MHC class I and II genes, suggesting increased antigen presentation activity. Additionally, we observed metabolic reprogramming in disease samples through increased expression of mitochondrial ATP synthesis genes and enrichment of oxidative phosphorylation and purine metabolism pathways. These findings demonstrate an interplay between neuroimmune activation and metabolic shifts in the SCG during cardiac disease. This re-analysis shows the value of revisiting existing transcriptomic datasets to uncover new biological insights.

## Introduction

The superior cervical ganglion (SCG) is a key component of the sympathetic nervous system, where it plays a role in the autonomic regulation of immune, metabolic, and cardiovascular functions. While sympathetic overactivation is a known feature of heart disease, the specific molecular and functional mechanisms that occur within the peripheral autonomic ganglia, such as the SCG, remain less clear.

A recent study by Ziegler et al. (1) profiled the SCG transcriptome in samples from both human cardiac disease patients and controls, in addition to corresponding mouse models using both bulk and single-cell RNA sequencing. While their main focus was on the single-cell data, the bulk RNAseq dataset remains a relatively underexplored resource. It holds the potential to reveal broader transcriptional programs involving immune activation, metabolic reprogramming, and neuroinflammation.

In this paper, we performed an independent reanalysis of the publicly available human SCG bulk RNA-seq data from Ziegler et al. (1) Our exploratory analysis uncovered transcriptional signatures that suggest immune cell recruitment, complement activation, antigen presentation, and metabolic shifts consistent with neuroinflammatory processes. These findings suggest that the SCG may serve as an active immuno-metabolic hub in the setting of cardiac disease and emphasize the value of revisiting transcriptomic datasets to gain new biological insights.

## Methods

### Dataset Acquisition

The bulk RNA-seq datasets analyzed in this study were obtained from the European Nucleotide Archive (ENA) under project accession number PRJNA967653. The dataset consists of six human superior cervical ganglia (SCG) samples, including three healthy samples and three samples from patients with cardiac disease.

### Data Preprocessing and Quality Control

Raw RNA-seq reads were downloaded to a Google Colab environment. Quality control was performed using FastQC (version 0.12.1) (2) on the Galaxy platform (usegalaxy.org). Trimming was not required, as quality metrics were within acceptable thresholds.

Reads were pseudoaligned and quantified at the transcript level using kallisto (version 0.46.2) (3), which was installed via conda (version 25.1.1) (Anaconda Software Distribution, https://www.anaconda.com). Pseudoalignment was performed against the human GRCh38 transcriptome (version 113).

The kallisto output was downloaded to a local machine and imported into R (version 4.4.2) using tximport (version (4). Gene annotations for GRCh38 (v113) were retrieved using biomaRt (version 2.62.1) (5, 6).

### Differential Expression Analysis

Low count genes (total counts <10 across all samples) were removed before analysis. Differential expression analysis was conducted using DESeq2 (version 1.46.0) (7). Genes were considered differentially expressed (DEGs) if they had an adjusted p-value < 0.05 and an absolute log2 fold change (|log2FC|) >= 1.

### Functional Enrichment Analysis

Gene Ontology (GO) and Kyoto Encyclopedia of Genes and Genomes (KEGG) enrichment analyses were performed using clusterProfiler (version 4.14.6) (8, 9) with annotations from org.Hs.eg.db (version 3.20.0) (10).

### Gene Set Enrichment Analysis

For Gene Set Enrichment Analysis (GSEA), genes were ranked based on log2 fold change. The msigdbr package (version 10.0.1) (11) was used to retrieve gene sets from the “C2” collection and “KEGG_LEGACY” subcollection. GSEA was done using fgsea (version 1.32.2) (12), setting a random seed of 123 for reproducibility.

### Data Visualization

Visualizations were produced using ggplot2 (version 3.5.1) (13), EnhancedVolcano (version 1.24.0) (14), pheatmap (version 1.0.12) (15), and enrichplot (version 1.26.6) (16).

### Data Handling and Manipulation

Data handling was performed using tidyverse (version 2.0.0) (17), dplyr (version 1.1.4) (18), readr (version 2.1.5) (19), and tibble (version 3.2.1) (20).

### Code and Reproducibility

All analysis code, Google Colab notebooks, R scripts, and FastQC files used in this study are openly available on GitHub at: (https://github.com/DjoserGenomics/SCG-cardiac-disease-reanalysis) to enable full reproducibility of the results.

## Results

### Quality Assessment and Sample Relationships

Principal Component Analysis (PCA) and a sample-to-sample distance heatmap were generated using variance-stabilized transformed (VST) counts to assess sample relationships and explore variation in gene expression.

PCA revealed two major sample clusters: CD01, CD03, and HS01 grouped on the negative side of PC1, while CD02, HS02, and HS03 clustered on the positive side (Figure 1). This indicates inter-sample heterogeneity and possible transcriptional similarity between some healthy and diseased samples. The sample-to-sample distance heatmap (Figure 2) supported this pattern and remained consistent after correcting for unwanted variation using Surrogate Variable Analysis (SVA), suggesting biological variability or a potential early disease state in CD03.

**Fig. 1.**
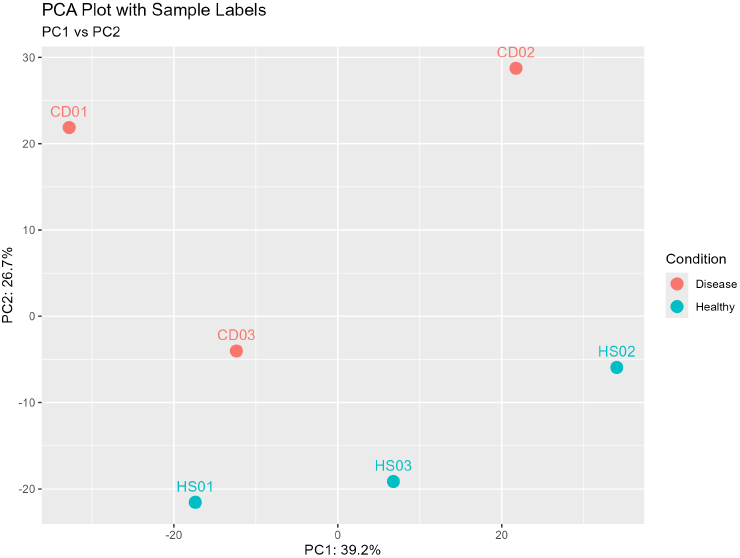
Principal Component Analysis of VST-transformed counts. CD01, CD03, and HS01 grouped on the negative side of PC1, while CD02, HS02, and HS03 clustered on the positive side.

**Fig. 2.**
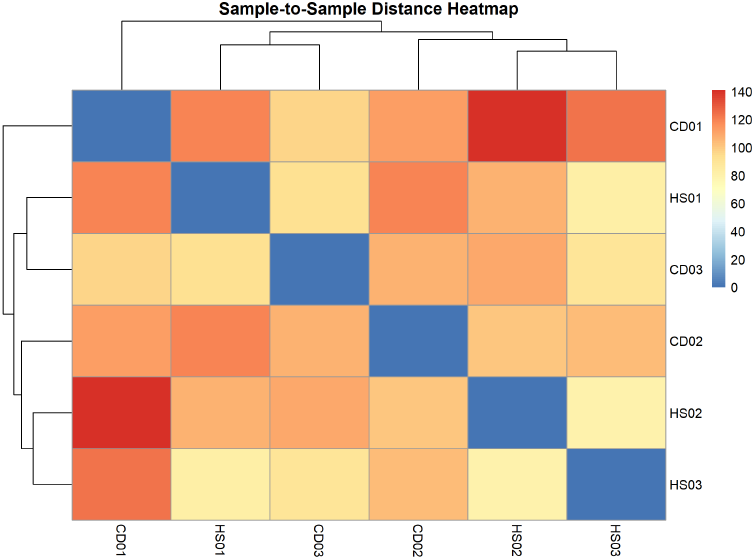
Sample-to-sample distance heatmap showing clustering of CD01, CD03, and HS01 together, and CD02, HS02, and HS03 as a second group.

Expression distribution boxplots (Figure 3) showed similar medians and spread across all samples, supporting the absence of technical biases or batch effects.

**Fig. 3.**
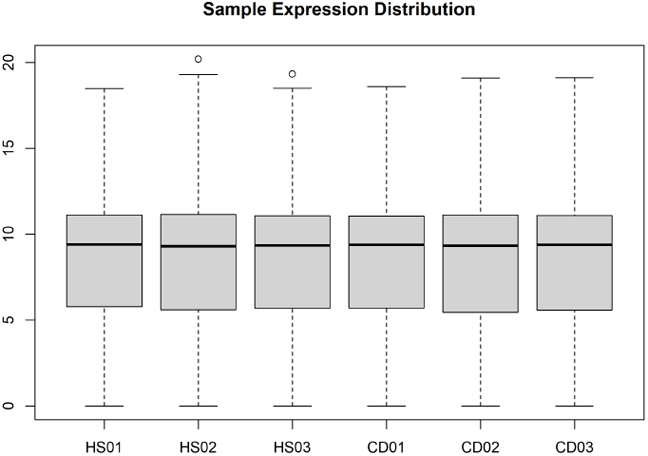
Expression distribution boxplots showing similar medians across samples, with only HS02 and HS03 having minor outliers.

### Differential Expression Analysis

Using DESeq2, we identified 771 differentially expressed genes (DEGs) with an adjusted p-value < 0.05 and (|log2FC|) >= 1. Among these, 399 genes were upregulated and 372 were downregulated in disease (CD) vs. healthy (HS) SCG samples. The full list of differentially expressed genes is available in Supplementary File 1.

The volcano plot (Figure 4) visualizes DEGs, with significantly altered genes highlighted in red. *DDTL* and *PTPRCAP* were strongly upregulated, while *NUDT4P2, ABCB11*, and *BIVM-ERCC5* were markedly downregulated, suggesting potential roles in disease pathology.

**Fig. 4.**
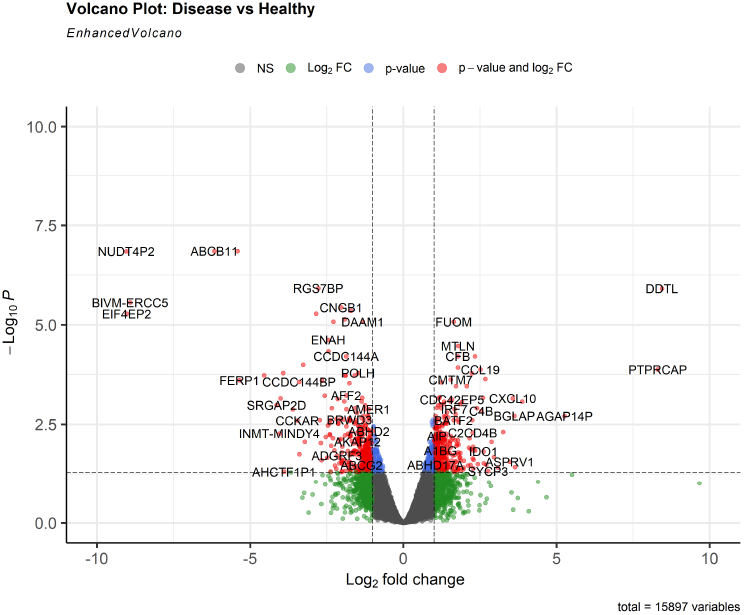
Volcano plot of DESeq2 results. Significantly differentially expressed genes (adjusted p<0.05 and |log2FC|>=1) are shown in red.

Neuroinflammation and immune activity emerged as prominent themes among the differentially expressed genes (DEGs). To visualize expression patterns of the associated genes, we generated two heatmaps.

The inflammatory cytokine heatmap (Figure 6) displays elevated or suppressed expression levels of selected pro-inflammatory markers across samples. Similarly, the immunity-related heatmap (Figure 7) highlights the differential expression of immune-related genes.

**Fig. 5.**
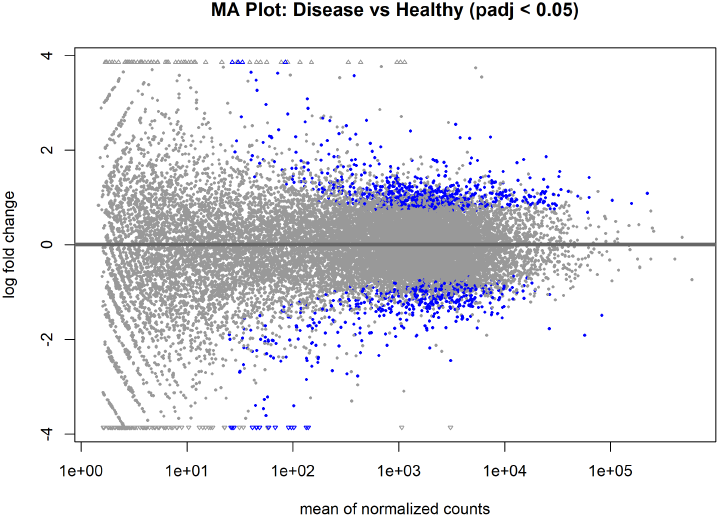
MA plot showing log2 fold change versus mean expression. Statistically significant DEGs are shown in blue. The symmetrical distribution confirms effective normalization.

**Fig. 6.**
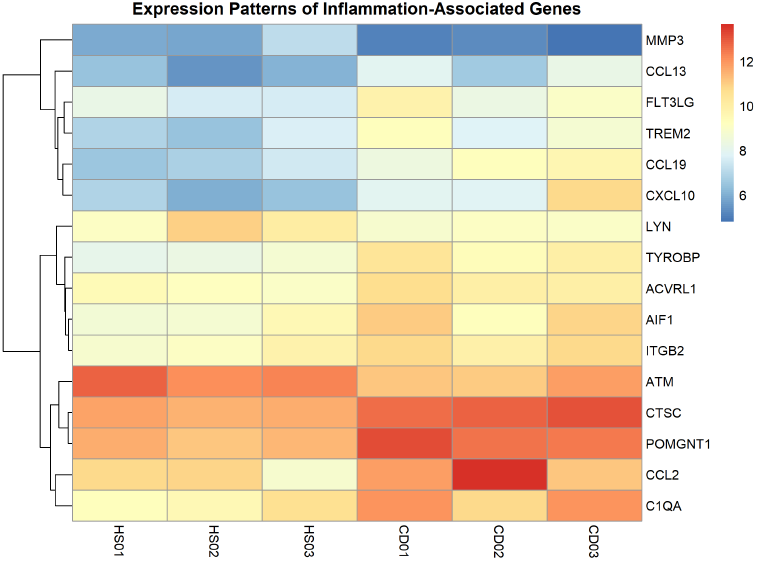
Heatmap showing expression levels of selected inflammatory cytokine genes across samples.

**Fig. 7.**
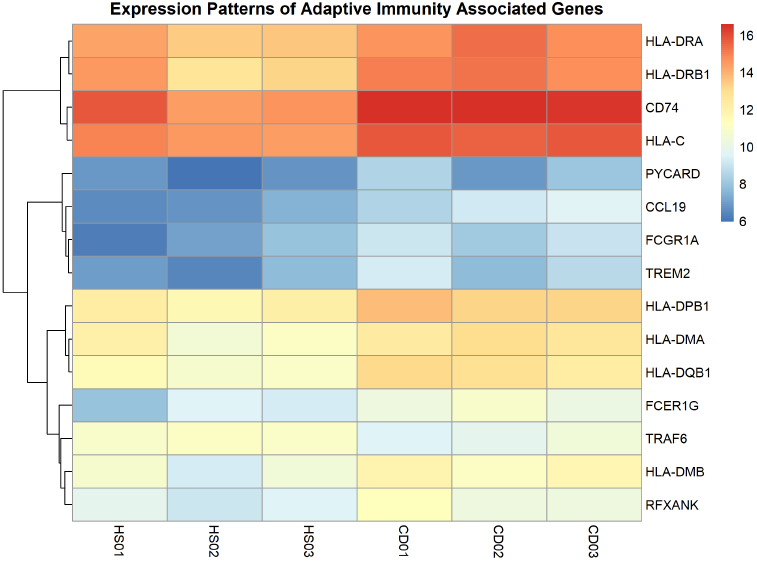
Heatmap showing expression of immunity-related genes across disease and healthy SCG samples.

### Functional Enrichment Analysis

GO enrichment identified 148 enriched terms (Figure 8): 119 biological processes (BP), 26 cellular components (CC), and 3 molecular functions (MF). Top BP terms included “Immune response-regulating cell surface receptor signaling pathway”, “Cytoplasmic translation”, and “Antigen processing and presentation via MHC class II”. All enriched GO terms are provided in Supplementary File 2.

**Fig. 8.**
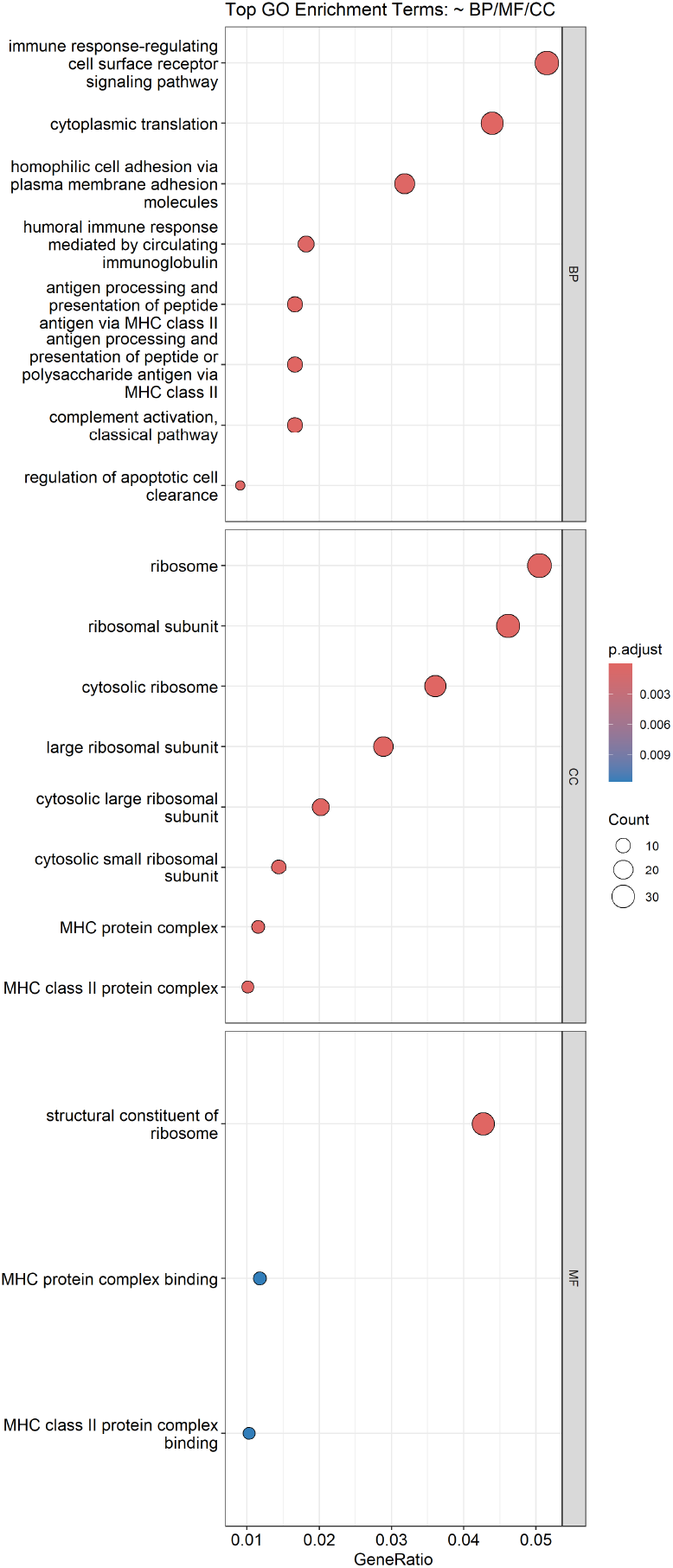
Gene Ontology (GO) Enrichment Dot Plot showing top Enriched terms across Biological Process, Cellular Component, and Molecular Function categories.

KEGG enrichment revealed 19 terms (Figure 9), including “Coronavirus disease – COVID-19”, “Staphylococcus aureus infection”, and “Pertussis”, likely reflecting shared immune/inflammatory mechanisms rather than specific pathology. All enriched KEGG pathways are provided in Supplementary File 3.

**Fig. 9.**
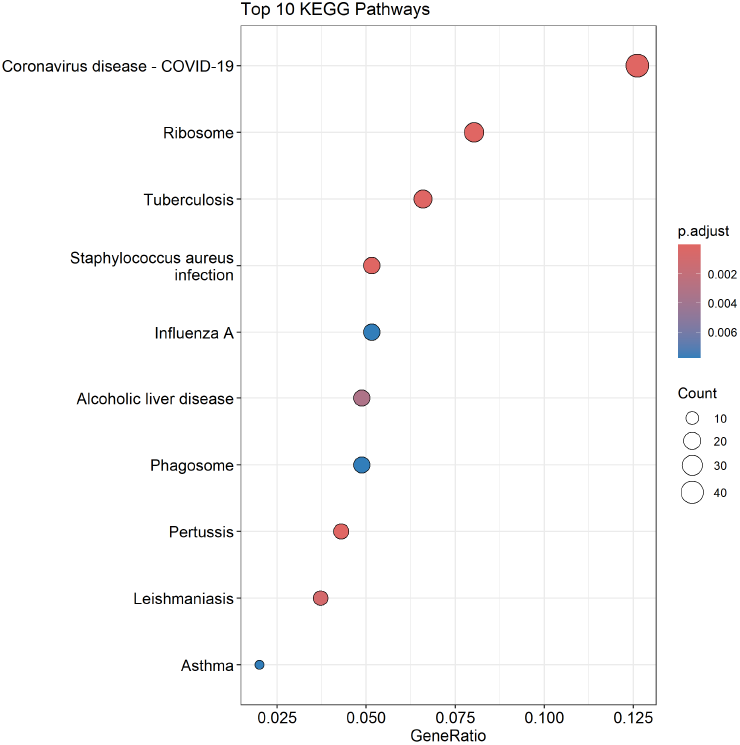
KEGG Enrichment Dot Plot showing Top enriched terms

### Gene Set Enrichment Analysis

Gene Set Enrichment Analysis (GSEA) identified 40 enriched gene sets, reinforcing immune and inflammatory themes. Enrichment Plots were generated for notably enriched sets, such as KEGG CYTOKINE CYTOKINE RECEPTOR INTERACTION (Figure 10) and KEGG ANTIGEN PROCESSING AND PRESENTATION (Figure 11). All enriched gene sets are available in Supplementary File 4. Alongside it, the ranked list of genes used to produce the GSEA results is available in Supplementary File 5.

**Fig. 10.**
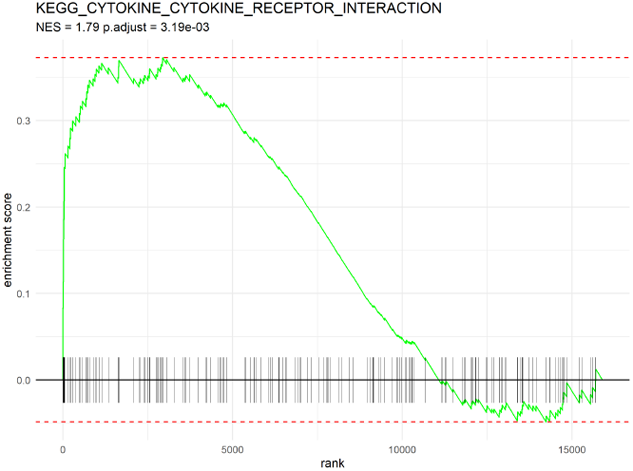
GSEA Enrichment plot of KEGG CYTOKINE CYTOKINE RECEPTOR INTERACTION shows initial spike followed by a decline suggesting an overall distribution

**Fig. 11.**
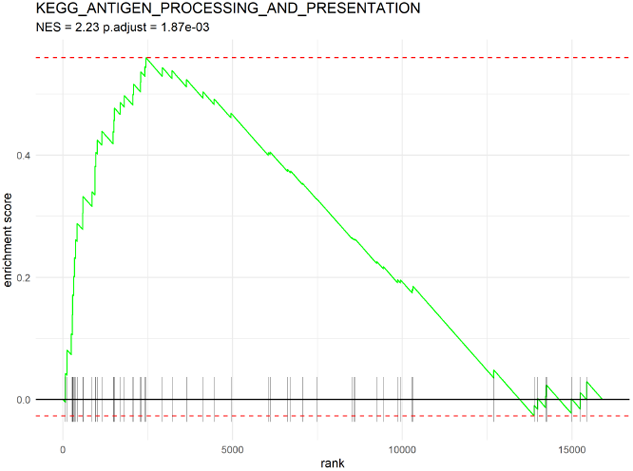
GSEA Enrichment plot of KEGG ANTIGEN PROCESSING AND PRESENTATION, it rises early and sharply, with a clear peak near the beginning

## Discussion

Neuroinflammation and immune activity were key responses in the superior cervical ganglia (SCG) of cardiac disease samples. This is supported by the upregulation of different chemokines such as CXCL10 (log2FC = 3.57, padj = 0.0007), CCL19 (log2FC = 2.51, padj = 0.0001) and CCL13 (log2FC = 2.28, padj = 0.0142), which indicates an activation of inflammatory cascades and immuno-regulatory responses. CXCL10 is known to attract T-cells and natural killer cells into neural tissues (21), CCL19 is important for leukocyte activation (22), and CCL13 attracts different kinds of immune cells and induces immunomodulatory responses (23). Together, they may be driving immune-mediated neuroinflammation in the SCG. This is further supported by the enrichment and upregulation of gene set enrichment analysis (GSEA) cytokine-cytokine receptor interaction pathway (NES = 1.79, padj = 0.0032).

Increased expression of TREM2 (log2FC = 1.86, padj = 0.0214) and TYROBP (log2FC = 1.70, padj = 0.0017) indicates activation of macrophage immune responses (24). This supports the macrophage infiltration in the SCG in cardiac disease samples reported by the original study through histological evidence (1). Gene Ontology (GO) analysis revealed enrichment of terms such as neuroinflammatory response (GO:0150076) and macrophage activation (GO:0042116), which provides additional support for the activated macrophage (1).

The enrichment of the KEGG Complement and Coagulation Cascades pathway (padj = 0.018) and differential expression of its associated genes, such as CFB (log2FC = 1.77, padj = 0.00006), C4B (log2FC = 2.54, padj = 0.0014), and C3 (log2FC = 1.73, padj = 0.0375), show increased evidence of complement-mediated activation of neuroinflammatory cascades and cytokine release (25) within the SCG during cardiac disease. KEGG Pathways associated with infectious or neurodegenerative diseases (e.g., hsa05171, hsa05322) were enriched, possibly due to shared immune and inflammatory actions, rather than disease pathology.

Beyond immune-related gene expression, enrichment of several Gene Ontology (GO) Biological Process terms pointed to altered purine nucleotide biosynthesis and metabolism, including purine nucleoside triphosphate biosynthetic process (GO:0009145), purine nucleotide biosynthetic process (GO:0006164), and purine nucleotide metabolic process (GO:0006163). These terms were driven by differential expression of mitochondrial complex I and ATP synthase genes, including NDUFS6 (log2FC = 1.22, padj = 0.0067), NDUFB11 (log2FC = 1.06, padj = 0.035), and ATP5F1D (log2FC = 1.32, padj = 0.0109), suggesting an increased need for energy in the SCG during cardiac disease. This aligns with the metabolic reprogramming seen in activated immune cells, such as macrophages, where M1-like activation shifts toward glycolysis and M2-like activation favors OXPHOS and fatty acid oxidation (26). Moreover, TREM2 provides a connection between purine metabolism and immune function and acts as an energy metabolism regulator in the peripheral nervous system (27). Which supports the link between metabolic and inflammatory processes in the SCG.

Furthermore, ATP can be produced through glycolysis in the cytosol or through oxidative phosphorylation in the mitochondria (28). Cardiac disease SCG samples showed increased oxidative phosphorylation through enrichment of the GSEA KEGG Oxidative Phosphorylation pathway (NES = 2.43, padj = 0.0018), suggesting a shift to mitochondrial ATP production in SCG cells during cardiac disease.

Antigen processing and presentation are key steps in initiating adaptive immune responses, which enable antigen-presenting cells (APCs) to display peptides to CD8+ (cytotoxic) and CD4+ (helper) T cells. GSEA revealed enrichment of the KEGG Antigen Processing and Presentation pathway (NES = 2.23, padj = 0.0019) in cardiac disease SCG samples, suggesting activation of adaptive immunity. Differential expression provides more evidence through upregulation of multiple genes encoding MHC class I and II molecules (e.g., HLA-C, HLA-DMA, HLA-DMB, HLA-DPB1, HLA-DQB1, HLA-DRA, HLA-DRB1).

These findings indicate an ongoing immune response in the SCG, potentially driven by antigenic stimuli. MHC class I molecules present peptides to CD8+ cytotoxic T cells, while MHC class II molecules present to CD4+ helper T cells, creating a coordinated adaptive response (29, 30). This adaptive immune activation may represent an immune reaction to neuroinflammatory changes in the SCG during cardiac disease.

## Limitations

A primary limitation of this study is the small sample size (n=3 per group), which can limit the statistical power to detect subtle transcriptomic changes and constrain the generalizability of the findings. Additionally, principal component analysis (PCA) revealed some heterogeneity among samples as discussed, suggesting possible biological variability or differing disease states. The transcriptomic changes identified here should be interpreted as exploratory and require validation in larger datasets and, ideally, through experimental approaches such as quantitative PCR or protein-level assays. However, this reanalysis offers valuable insights into the molecular and functional mechanisms of the SCG in cardiac disease and lays a foundation for future investigations.

## Conclusion

This transcriptomic reanalysis of the superior cervical ganglia (SCG) in cardiac disease highlights a coordinated immune and inflammatory response. We identified upregulation of inflammatory chemokines, activation of macrophage-related genes, and enrichment of antigen presentation pathways, all pointing to neuroinflammation and adaptive immune activation in the SCG. Additionally, enhanced expression of mitochondrial and purine metabolism genes, alongside oxidative phosphorylation pathway enrichment, suggests a metabolic change to support immune cell function. Together, these findings support a model in which cardiac disease drives immune-mediated remodeling and metabolic reprogramming in the SCG.

## Supporting information

Supplementary Table 1 - Differentially Expressed Genes (DEGs)

Supplementary Table 2 - Gene Ontology (GO) Enrichment Analysis Results

Supplementary Table 3 - KEGG Enrichment Analysis Results

Supplementary Table 4 - GSEA Results

Supplementary Table 5 - Ranked Genes List for GSEA

R Session Info

